# Development of loop-mediated isothermal amplification (LAMP) assay for detection of Pseudocercospora angolensis in sweet orange

**DOI:** 10.1101/2021.01.13.426516

**Authors:** Patricia Driciru, M Claire Mugasa, Robert Acidri, John Adriko

## Abstract

*Pseudocercospora angolensis* is the causative agent of *Pseudocercospora* leaf and fruit spot disease in citrus which can result in up to 100% yield loss. Early diagnosis of this disease is vital for effective control. This study aimed at developing a loop-mediated amplification (LAMP) system for detecting *P. angolensis* in sweet oranges in comparison with Polymerase Chain Reaction (PCR) and using microscopy as a gold standard. Twelve non-target species were used to assess the analytical specificity of LAMP and PCR whereas the analytical sensitivity was determined using serial dilutions of *P. angolensis* DNA. The diagnostic accuracies of the two assays were evaluated using DNA from 150 diseased and 50 non-diseased sweet orange leaf samples. The analytical sensitivity and detection time of LAMP were of 10^−4^ ng/ μl and 40 minutes, respectively. The analytical sensitivity of PCR was 10ng/μl and it was specific to *P. angolensis* whereas three relatives of *P. angolensis* were detectable by LAMP. The diagnostic sensitivities of LAMP (93%) and microscopy (100%) were significantly different (*X*^*2*^ = 8.38, *P* = 0.0038) unlike the diagnostic specificities (90%) and (100%), respectively (*X*^*2*^ = 3.37, *P* = 0.066). Microscopy was significantly more sensitive than PCR (32.6%) (*X*^*2*^ = 149.26, *P* < 2.2e-16) and equally specific as PCR (*P*=NA). The positive predictive values of PCR and LAMP were 100% and 96.5% respectively whereas the negative predictive values were 33.1% and 81.8% respectively. The LAMP assay developed in this study offers a great tool for routine screening sweet orange samples for *P. angolensis*.

## 1. Introduction

Sweet orange (*Citrus cinensis*) is the second most widely grown fruit crop in the world by volume next to banana [1]. *Pseudocercospora* leaf and fruit spot disease, caused by *Pseudocercospora angolensis*, also known as angular leaf spot disease, is one of the main challenges constraining lucrative production of citrus in Africa [2, 3]. This disease can cause 50 to 100% yield loss in affected orchards [4,5]. Early, rapid, accurate, fast and safe detection is required for effective control of the disease. The symptoms of *Pseudocercospora* leaf and fruit spot can be confused with those of citrus canker and citrus black spot especially during early pathogenesis [6, 2]. Traditional methods of plant pathogen detection are time consuming and tedious [7]. Molecular methods such as PCR have been used to detect *P. angolensis* [8, 9]. Despite the specificity and sensitivity of PCR, this method is expensive due to the cost of thermocyclers and consumables used, hence not very cost-effective in low-income nation settings [10]. Thermocyclers might also be erroneous at high ambient temperatures, humidity and dusty environments [11]. Results of PCR may only be obtained after 2-3 hours, which makes it a slow technique [11]. Thus, there is need for a simple, cost-effective, and rapid DNA amplification method for diagnosis and monitoring of *P. angolensis* for effective management of the disease. Loop-mediated isothermal amplification of DNA (LAMP), a method developed by [12] has become an alternative to conventional PCR. Amplification and detection occur in a one-step reaction in a single tube [13]. LAMP is a highly sensitive, efficient, cheaper, and simple DNA amplification technique applicable in pathogen screening in germplasm in low-income settings. LAMP has already been developed for several bacteria pathogens namely, *Erwinia amylovora* in pear and apple, *Ralstonia solanacearum* in tomatoes, *Napier stunt phytoplasma* in Napier grass and *Xanthomonas arboricola* in *Prunus spp*. and viruses such as potato virus Y [14, 15, 16, 17, 18]. Very few LAMP detection assays have already been optimized for some fungi such as *Fusarium graminearum* in wheat and wheat stripe rust but this method has not been tested for citrus fungal pathogens including *P. angolensis* [19, 20]. Thus, the purpose of this study was to develop LAMP detection of *P. angolensis* to enhance easy detection and management of *Pseudocercospora* leaf and fruit spot disease in citrus.

## 2. Materials and methods

### 2.1 Microorganism sourcing, sample collection and preservation

Reference gDNA samples of *Pseudocercospora angolensis* (LSVM 1309, LSVM 1308, and MC 39) and non-targets namely *P. fijiensis* (LSVM501)*, P. musicola* (LSVM 511) and *P. eumusae* (LSVM 536) were obtained from Agence Nationale de Securite Sanitaire de l’Alimentation, de l’Environnement et du Travail (ANSES), France, whereas the reference gDNA of *P. griseola* and pure culture of *Colletotrichum lindemuthanium* were sourced from International Center for Tropical Agriculture (CIAT), Uganda. Citrus pathogens namely *Guignardia citricarpa*, *Pseudocercospora angolensis*, *Colletotricum gleosporoides* were isolated from lesions on sweet orange leaf and fruit samples and confirmed by microscopy. On the other hand, the gDNA of other citrus pathogens including *Aternaria alternata*, *Elsinoe Fawcetti, Diaporthe citri* and *Phytophthora citrophthora* were directly extracted from lesions on sweet orange fruit and bark (for *Phytophthora citrophthora* only) samples were collected from Bukedea district in Eastern Uganda and each sample placed in a separate plastic bag to avoid cross contamination. Additionally, hands were disinfected with 70% ethanol after collection of each sample. These samples were transported to the National Agricultural Research Laboratories (NARL) for analysis.

### 2.2 Pathogen isolation and DNA extraction

Sweet orange fruit samples symptomatic for *Pseudocercospora* leaf and fruit spot, citrus black spot and anthracnose were used to isolate *Pseudocercospora angolensis*, *Guignardia citricarpa* and *Colletotrichum gleosporoides*, respectively. The samples were washed under running tap water with liquid soap and surface sterilized in 70% ethanol for 5 minutes, rinsed with sterile water and surface sterilized again in 5% Sodium hypochlorite for 1 minute and thoroughly rinsed in three changes of sterile water. Three small pieces were removed from area around a lesion per fruit sample with a sterile scalpel blade and inoculated on Potato Dextrose Agar. Six replicates of the cultures were made and incubated at 25±1°C. Five days after inoculation, the cultures were examined for growth of the above three citrus pathogens. Colonies with the expected morphological features were sub-cultured on fresh potato dextrose agar (PDA), oatmeal agar (OA) and V8 Juice media. The cultures were then incubated for 14 days at 25±1°C while exposing them to 13h of natural light and 11 h of fluorescent light. Each isolate was identified based on the phenotypic colony morphologies and using a light microscope (Karl Zeiss) to visualize the hyphae and conidial structures of each of the pathogens.

Genomic DNA of *P. angolensis*, *G. citricarpa*, *C. gleosporoides* and *C. lindemuthanium* were extracted from the mycelia whereas those of *A. alternata*, *E. Fawcetti, D. citri* and *P. citrophthora* were directly extracted from symptomatic sweet orange fruit and bark samples (in the case of *P. citrophthora* only). Extraction of DNA was done using the Cetyl Trimethyl Ammonium Bromide (CTAB) method and the eluted DNA quantified using nanodrop 2000c (Thermoscientific).

### 2.3 *In-silico* primer design

A PCR primer set designed by Ahmed *et al*., (2018) (Ango-Conv) was used to confirm *P. angolensis* isolates in this study. A novel primer pair (Cer 2) was designed for optimization to target the small subunit rRNA gene, 5.8S, intergenic regions, ITS1, and ITS2, and partial sequence of the large subunit ribosomal RNA in the genome of *P. angolensis* with a GeneBank accession number MH857140.1. The uniqueness of the marker sequence was verified through an nBLAST and multiple sequence alignment. On the other hand, four primer sets were designed for the LAMP assay to target six regions on the template and these primers included the forward outer primer (F3), forward inner primer (FIP), backward outer primer (B3) and backward inner (BIP) primer, loop forward primer (LF) and loop reverse primer (LR). The LAMP primers were designed based on the internal transcribed spacer (ITS) region and the translation elongation factor 1 alpha (*Tef* 1 alpha) gene of *Pseudocercospora angolensis* with NCBI GenBank accession numbers MG956768.1 and JX901668.1, respectively using primer explorer V5 software. The specificity of the primers was ascertained by performing a BLAST search against available sequences in the GenBank. The quality of the primers was analysed using IDT oligo analyser for potential secondary structure formation, complimentary palindromes, and primer dimer formation within and between different primers. Following *in-silico* analysis of specificity of the primer sets, one ITS-based primer set (ID1-ITS) and one *Tef*-based primer set (ID1-TEF) were chosen for optimization. Details of the primers are presented in Table 1 below.

**Table 1.**
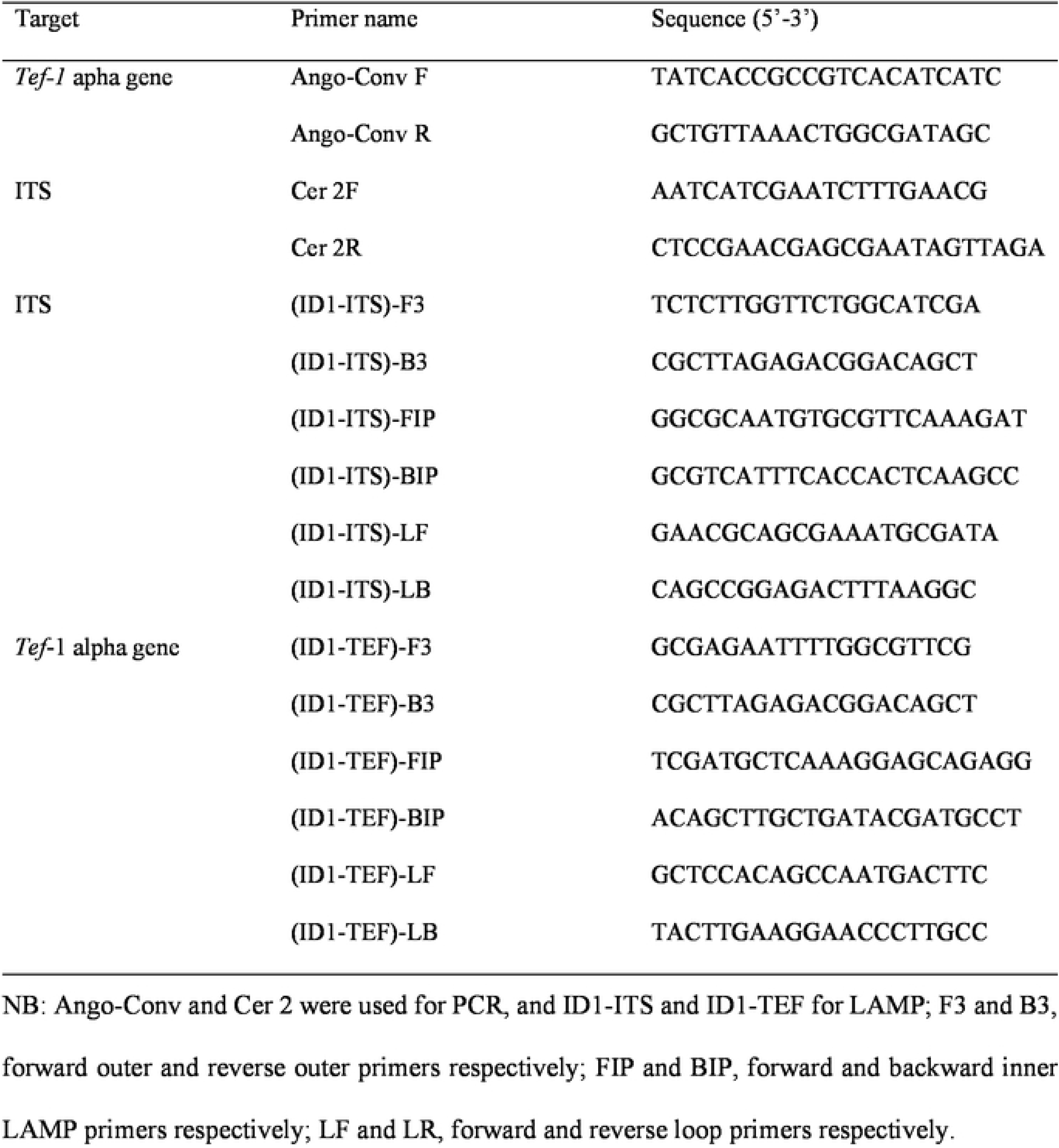
Primers used in this study.

**Table 2.** Proportions of true positive, false positive, true negative and false negative values obtained in PCR detection of *P. angolensis* in sweet orange leaf samples using the Ccr 2 primers.

| **PCR** | **Microscopy** |  |
| --- | --- | --- |
|  | Positive | Negative |
| Positive | 49 (TP) | 0 (FP) |
| Negative | 101 (FN) | 50 (TN) |

NB: TP: True positive, FP: False positive, TN: True negative, FN: False negative

**Table 3.** Proportions of true positive, false positive, true negative and false negative values obtained in LAMP detection of *P angolensis* in sweet orange leaf samples

| **LAMP** | **Microscopy** |  |
| --- | --- | --- |
|  | Positive | Negative |
| Positive | 140 (TP) | 5 (FP) |
| Negative | 10 (FN) | 45 (TN) |

NB: TP: True positive, FP: False positive, TN: True negative, FN: False negative

### 2.4 PCR Optimization for detection of *P. angolensis*

PCR was performed in a 20 μl reaction using the Accupower® TaqPreMix (0.2ml/50μl reaction) from Bioneer. Two (2) μl of the forward primer, 2 μl reverse primer (10μM), 2 μl DNA (25ng/μl) and 14 μl of water were added to the Taq premix and dissolved. The reaction components were then spun for two seconds using a rotor (Techino Mini). DNA amplification was carried out in a C 1000 Touch™ thermocycler (BIORAD). The reference gDNA samples of *P. angolensis* and gDNA purified from a pure culture of *P. angolensis* were used as templates to optimize PCR amplification. For PCR reactions where the ango-conv primer set was used and pre-optimized PCR conditions by Ahmed *et al*., (2018) were used. Reaction conditions for PCR were optimized for the Cer2 primer and these included: initial denaturation at 95°C for 10 min followed by 30 cycles of denaturation at 94°C for 30s, annealing at 45 to 49°C for 30s and extension at 72°C. Nuclease-free water was used as the internal negative control whereas gDNA of *Citrus cinensis* was used as the external negative control. The PCR components included template DNA, dNTPs, *Taq* polymerase, primers, Magnesium chloride, and buffer. The PCR products (5μl) were separated on 1.5% agarose gel in 1X TBE buffer by gel electrophoresis, and the bands were stained using Ethidium bromide and visualized under UV light.

### 2.5 Optimization of LAMP for *P. angolensis* detection

Conditions for detecting *P. angolensis* by LAMP assay were standardized using the following reaction components: DNA template, 25 mM Magnesium chloride (0.4-0.8 μl), 10mM dNTPs (0.8μl), primers F3 and B3 (1-3μM), FIP and BIP (0.5-3μM), LF and LB (0.1-1μM), 0.32 units/ μl sapphire *Bst* 2.0 polymerase (Jena Bioscience), *Bst* 2.0 polymerase buffer (2 μl) (Jena Bioscience). The incubation temperature was varied from 60-65°C for 60 min. The minimum detection time for the LAMP assay was determined through detecting *P. angolensis* using the optimized LAMP reaction components at different durations (0 min, 10 min, 20 min, 30 min, 40 min, 50 min and 60 min) of incubation of the LAMP reaction components. Both the ID1-ITS and ID1-TEF primers were optimized at the above conditions. The LAMP products (3 μl) were separated through 2% agarose in 1X TBE at 100v for 60-90 minutes, stained in Ethidium bromide and viewed under UV.

### 2.6 Determination of the analytical accuracy of PCR and LAMP assays

The analytical sensitivity of PCR was assessed with serial dilutions (1000ng/μl-10^−4^ ng/μl) of *P. angolensis* gDNA positive control. To determine the analytical sensitivity of LAMP, ten-fold serial dilutions (100 ng/μl-10^−5^ng/μl) of *P. angolensis* DNA were prepared. The limit of detection was determined as the minimum quantity of target DNA that could consistently be amplified by PCR and LAMP. Reproducibility was tested with one replicate of the same DNA concentration during an individual run. Each run was repeated twice and in case of a disagreement in the assay results, it was repeated for the third time for a more conclusive result. The analytical sensitivity of the assays was determined through obtaining the minimum detection limits of either assay.

The analytical specificity involved testing the ability of the two assays to detect only *Pseudocercospora angolensis* and not the non-targets namely *Pseudocercospora emusae*, *Pseudocercospora musicola*, *Pseudocercospora fijiensis*, *Guignardia citricarpa*, *Colletotrichum gleosporoides*, *Colletotrichum lindemuthianum*, *Aternaria alternata*, *Elsinoe fawcetti, Diaporthe citri*, *Phytophthora citrophthora* and *Citrus cinensis*. All DNA was standardized to the lowest detection limits of the assaysand DNA from healthy orange leaf tissue was used as the external negative control whereas nuclease-free water was used as the internal negative control. The analytical accuracy of both the ID1-ITS and ID1-TEF primer sets were determined so that the primer set with a better analytical specificity and sensitivity would be chosen to evaluate the diagnostic accuracy of LAMP in the detection of *P. angolensis*.

### 2.7 Inoculation of sweet orange plants, leaf sample collection and DNA extraction

A total of 200 healthy Washington navel plants were obtained from the National Agricultural Research Laboratories (NARL) nursery and pathogenicity test was performed once on healthy 6-month-old Washington navel plants and these plants were divided into two groups: one group consisting of 150 plants maintained in one compartment of the screen house and the other group included 50 plants in a separate compartment. The 150 plants were inoculated with conidia (unquantified due to conidia being sparse) by spraying the leaves to run off, placed in a humid chamber and allowed to develop symptoms of *Pseudocercospora* leaf spot. The other group consisting of 50 plants was not inoculated. Genomic DNA was extracted from 150 symptomatic leaf samples four weeks after the onset of symptoms of *Pseudocercospora* leaf spot and 50 asymptomatic leaf samples using the CTAB method.

### 2.8 Determination of the diagnostic accuracy of PCR and LAMP assays

The optimized LAMP and PCR assays were then used to detect *P. angolensis* in the DNA samples extracted from the sweet orange leaves with known infection status in the two groups mentioned above and detected by LAMP and PCR. On the other hand, *P. angolensis* was isolated from the lesions of the symptomatic leaf samples and the conidia were identified by a light microscope (Carl Zeiss) (x40). Detection of samples by each assay was repeated twice and in any case of disagreement, it was repeated for the third time. True positive, true negative, false positive and false negative values were then recorded for each assay. These values were used to calculate the sensitivity, specificity, positive predictive value and negative predictive value of PCR and LAMP using statistical formulae by [21] below.

Sensitivity (SE)= true positive (TP)/ (true positive (TP) + false negative (FN))*100% Specificity (SP) = true negative (TN)/(true negative (TN) + false positive (FP))*100% Positive predictive value (PPV)= true positive (TP)/ (true positive (TP)+ false positive (FP))*100% Negative predictive value (NPV) = true negative (TN)/ (true negative (TN) + false negative (FN))*100%.

### 2.9 Statistical analyses

The data generated was entered in Microsoft Excel (2010) and summarized in contingency tables. The sensitivity, specificity, positive and negative predictive values of the two assays was then calculated independently using statistical formulae described by [21]. R statistical software; RStudio-0.98.1102 was used to analyze the data. The sensitivity and specificity of PCR and LAMP were each compared with microscopy using Pearson’s chi-square test to evaluate the diagnostic accuracy of each assay.

## 3. Results

### 3.1 Isolation of *Pseudocercospora angolensis*,*Guignardia citricarpa* and *Colletotricum gleosporoides*

*P. angolensis* had slow-growing, grey colonies with a dark green color on the underside (fig 1a). The conidia were arranged in chains (white arrow in figs 1 b & c), with 3 to 6 conidia per chain. The colonies of *G. citricarpa* were black and compact with invaginations surrounded by a yellow pigment (fig 1d) whereas the conidia were oval (figs 1e & 1f). *C. gleosporoides* mycelia were white at first but later turned grey with some orange pigmentation (fig 4g). The conidia were cylindrical with rounded apices (fig 1h).

**Fig 1.**
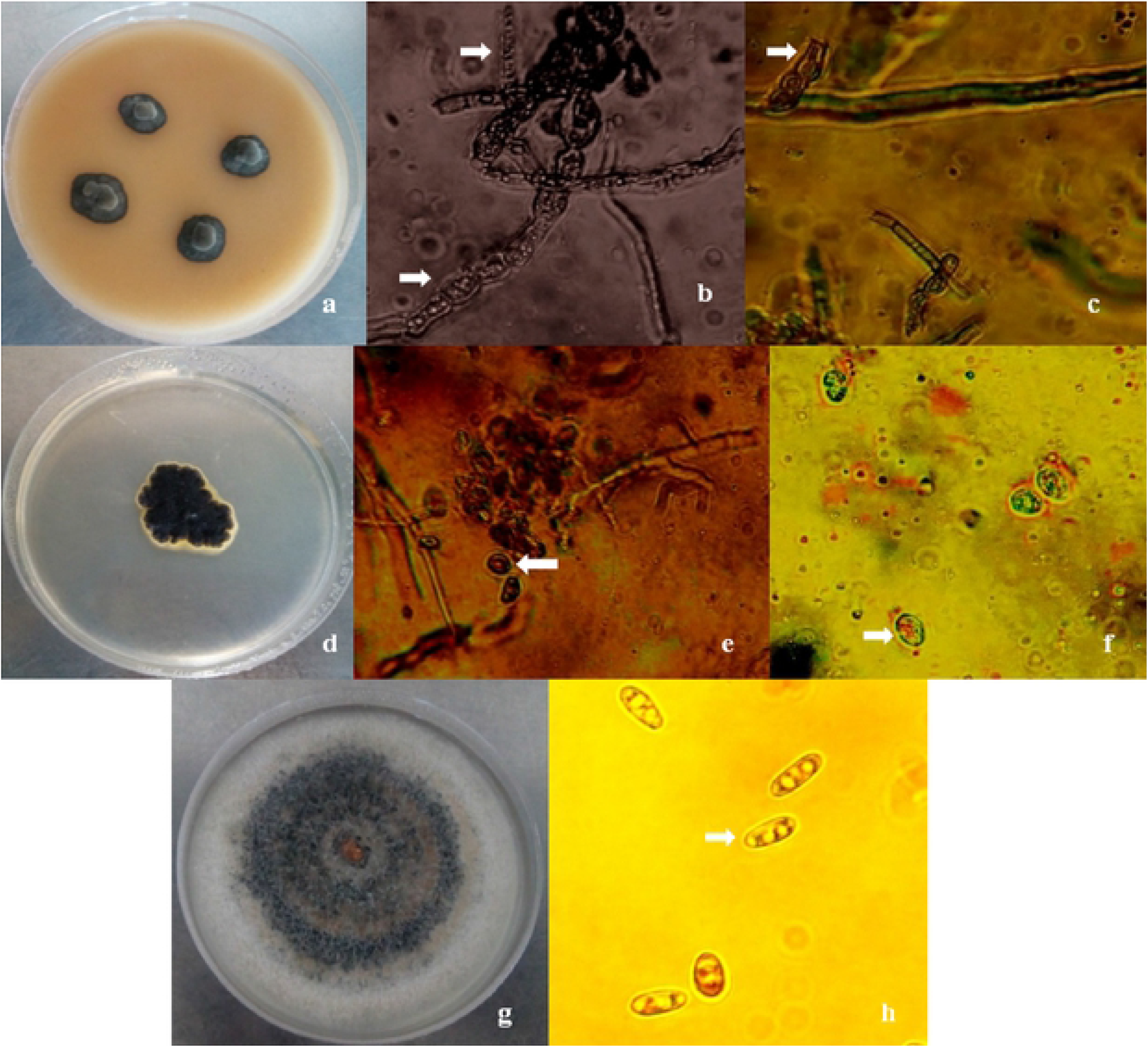
Cultures and conidia of *P. angolensis, G. citricarpa* and C. *gleoporoides;* a- colonies of *P. angolensis* on V8, b & c- *P. angolensis* conidia (see arrow), d- colony of *G. citricarpa* on PDA, e & f- conidia of *G. citricarpa* (see arrow), g- colony of *C. gleosporoides* on oatmeal agar h- *C. gleosporoides* conidia (see arrow).

### 3.2 Optimization of PCR for detection of *P. angolensis*

All the reference *P. angolensis* gDNA samples and the gDNA sample extracted from an isolate of *P. angolensis* were amplified at optimum PCR conditions for each of the two PCR primer pairs used (Cer2 and Ango-Conv). Primer pair Cer 2 produced a band of 188bp, whereas the primers designed by [8] (Ango-Conv) produced a band of 218 bp as shown in fig 2. The optimum PCR conditions for the Cer 2 primer included initial denaturation 95°C for 10 min, second denaturation at 94°C for 30s, annealing at 49°C for 30s extension at 72°C for 30s and final extension at 5 min.

**Fig 2.**
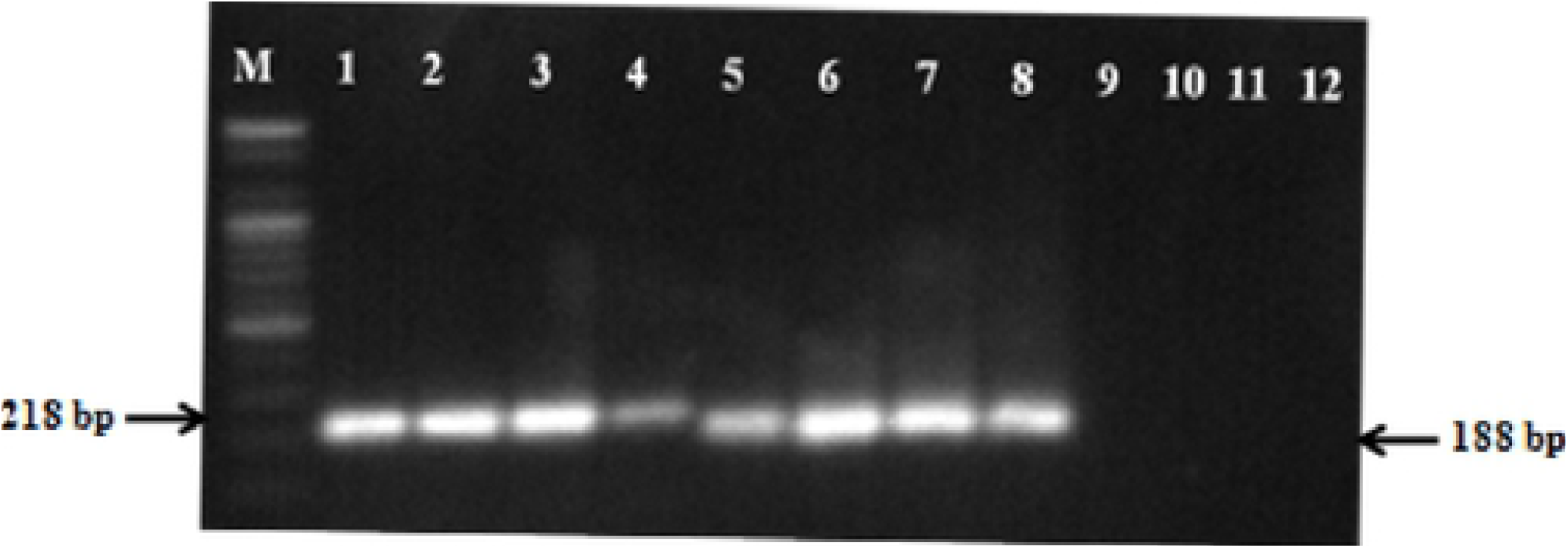
Amplification of *P. angolensis* gDNA using the standardized PCR protocol for the Ango-Conv primers (samples 1-4) and Cer2 primers (samples 5-8); M-100bp ladder 1 & 5- gDNA from the mycelia of *P. angolensis* isolated from a fruit sample, 2 & 6- *P. angolensis* strain LSVM 1309, 3 & 7- *P. angolensis* strain LSVM 1308, 4 & 8- *P. angolensis* strain MC 39, 9 & 11- external control *(Citrus cinensis* gDNA) and 10 & 12- internal negative control (nucleic aci dfree water)

### 3.3 Determination of the analytical accuracy of PCR in *P. angolensis* detection

#### 3.3.1 Assessment of the analytical specificity of PCR in the detection of *P. angolensis* using Cer 2 primers

The PCR based marker showed high specificity whereby only the target DNA templates tested positive. Reactions where negative controls were used as template also did not show any amplification as shown in fig 3.

**Fig 3.**
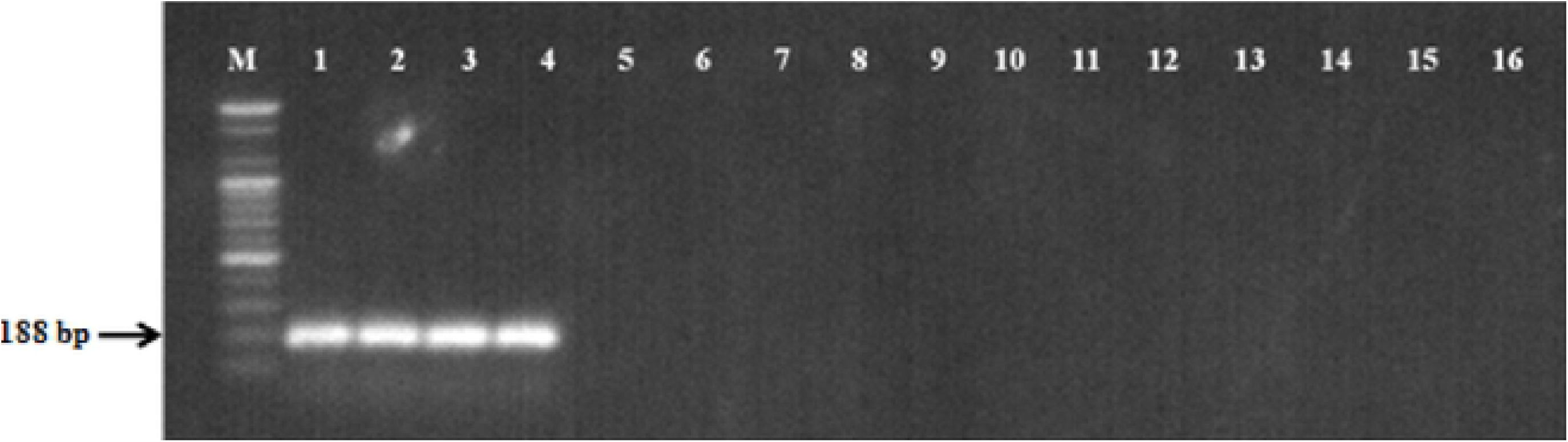
Shows specific PCR amplification of *P. angolensis* DNA: M- 100bp ladder, 1-DNA extracted from an isolate of *P. angolensis*, 2-4- Reference samples of *P. angolensis* (2- LSVM 1309, 3- LSVM 1308, 4- MC 39), 5- *P. griseola*, 6- *P. emusae*, 7- *P. musicola*,8- *P. fijiensis*,9- *C. gleosporoides*, 10- *E. Fawcelli*, 11- *G. citricarpa*, 12- *A.altemata*, 13-*C. lindemuthianum*, 14- *D. citri*, 15- *P. citrophthora*, 16- Negative control (citrusDNA).

**Fig 4.**
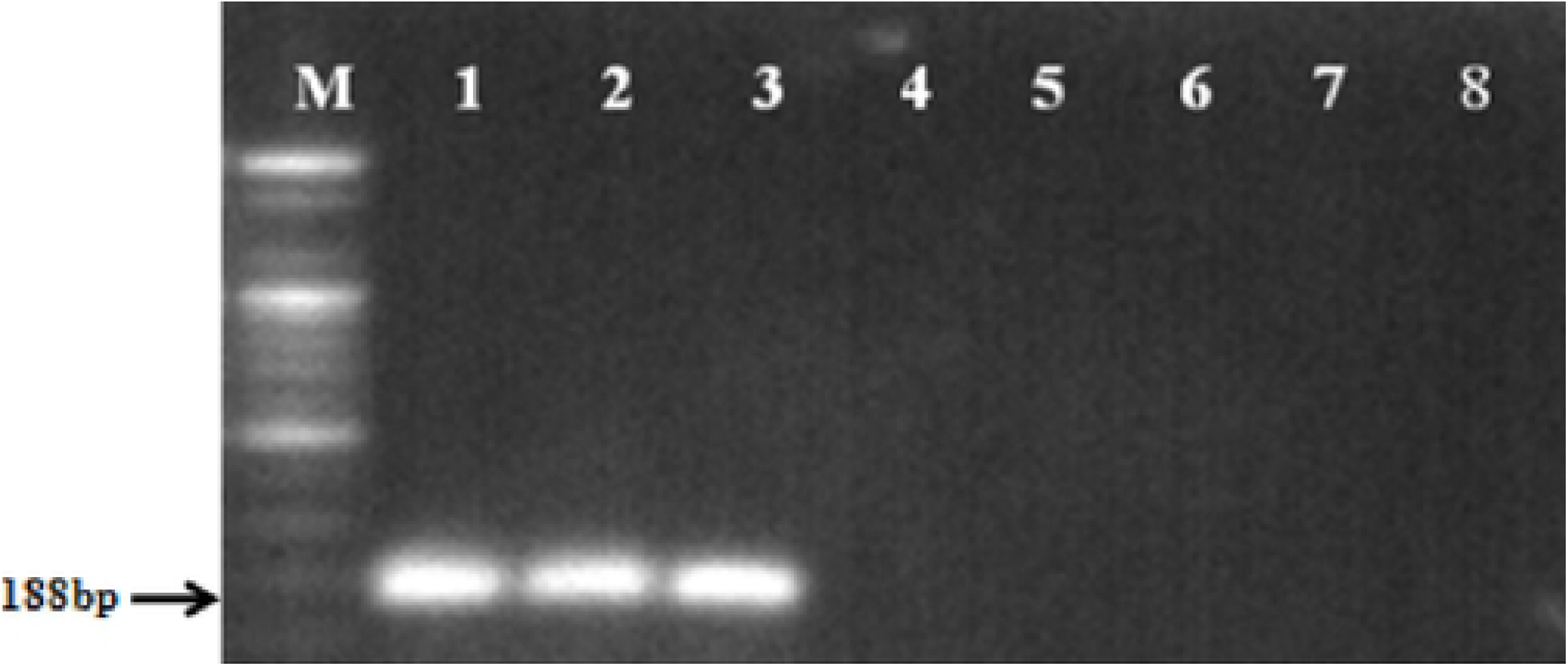
Sensitivity of PCR in detecting *P. angolensis* using the Cer2 primers; M-100 bp ladder, 1-1000ng/μl, 2-100ng/μl, 3-10ng/μl, 4-lng/μl, 5-10^−1^ng/μl, 6-10^−2^ng/μl, 7-10^−3^ng/μl, 8-10^−4^ng/μl.

#### 3.3.2 Assessment of the analytical sensitivity of PCR in *P. angolensis* detection using Cer 2 primers

Of the different concentrations that were detected, amplification was seen in PCR reactions that had *P. angolensis* gDNA template concentrations of 1000ng/μl, 100ng/μl and 10ng/μl. Template concentrations of 1 ng/μl, 10^−1^ ng/μl, 10^−2^ ng/μl, 10^−3^ ng/μl and 10^−4^ ng/μl were not amplified as illustrated in fig 4.

### 3.4 Optimization of LAMP for detection of *P. angolensis*

#### 3.4.1 Standardization of LAMP conditions using the ID1-ITS primer set

The optimized concentrations of the components in 20 μl reaction included 2 μl of the 10X sapphire *Bst* 2.0 polymerase buffer, 1mM MgCl_2_, 400μM dNTP mix, 0.32 units/μl sapphire *Bst* 2.0 polymerase, 2 μM of FIP and BIP, 0.25 μM of F3 and B3 and 1 μM of LF and LB primers. The optimum temperature of the reaction was 64°C (fig 5a) whereas the minimum time at which amplification was visible was at 20 minutes. At 30 minutes of incubation, the bands were clearer as shown in fig 5b.

**Fig 5.**
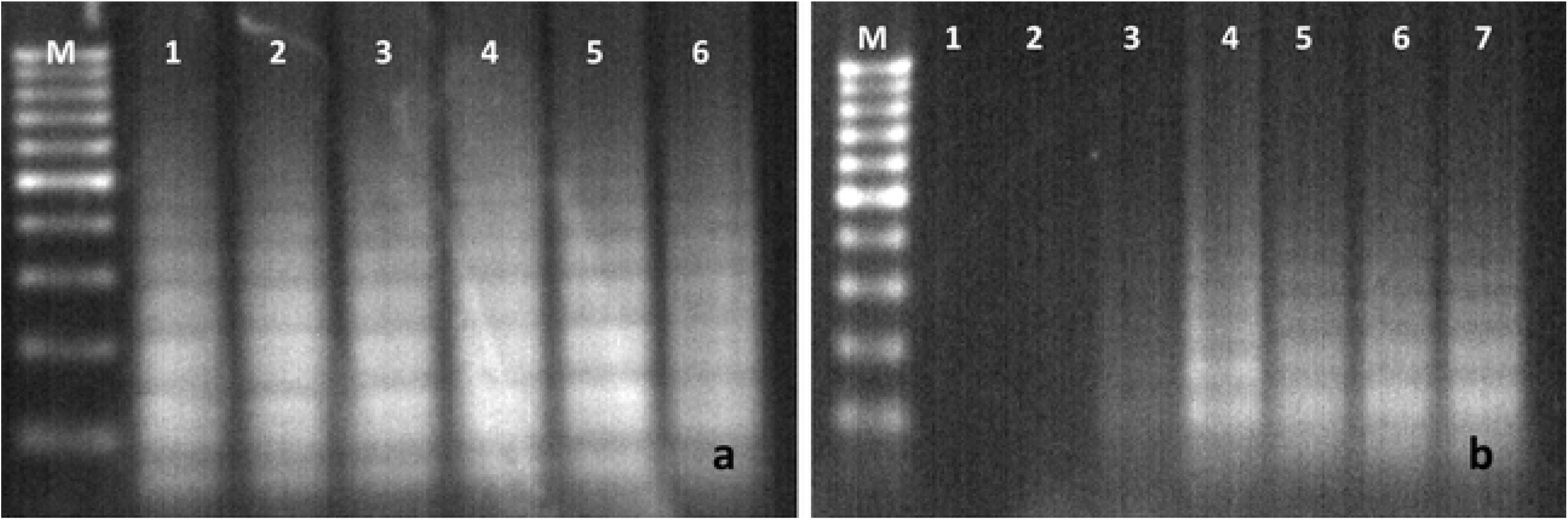
LAMP amplification of *P. angolensis* DNA (LSVM 1309) using ID1-ITS primers; a- at different temperatures; M-100bp ladder, 1-60 °C, 2-61 °C, 3-62 °C, 4-63 °C, 5-64°C, 6-65°C; b- at diferent incubation times; M-100bp ladder, 1-0 min, 2-10 min, 3-20 min, 4-30 min, 5-40 min, 6-50 min and 7-60 min

#### 3.4.2 Standardization of LAMP conditions using the ID1-TEF primer set

The optimum concentrations of the LAMP reaction components in 20 μl reaction included 2 μl of the 10X sapphire *Bst* 2.0 polymerase buffer, 1mM MgCl_2_, 400μM dNTP mix, 0.32 units/μl sapphire Bst 2.0 polymerase, 2 μM of FIP and BIP, 0.25 μM of F3 and B3 and 1 μM of LF and LB primers. The optimum temperature for the reaction was 65°C (fig 6a) while shortest time of detection was 40 min (fig 6b).

**Fig 6.**
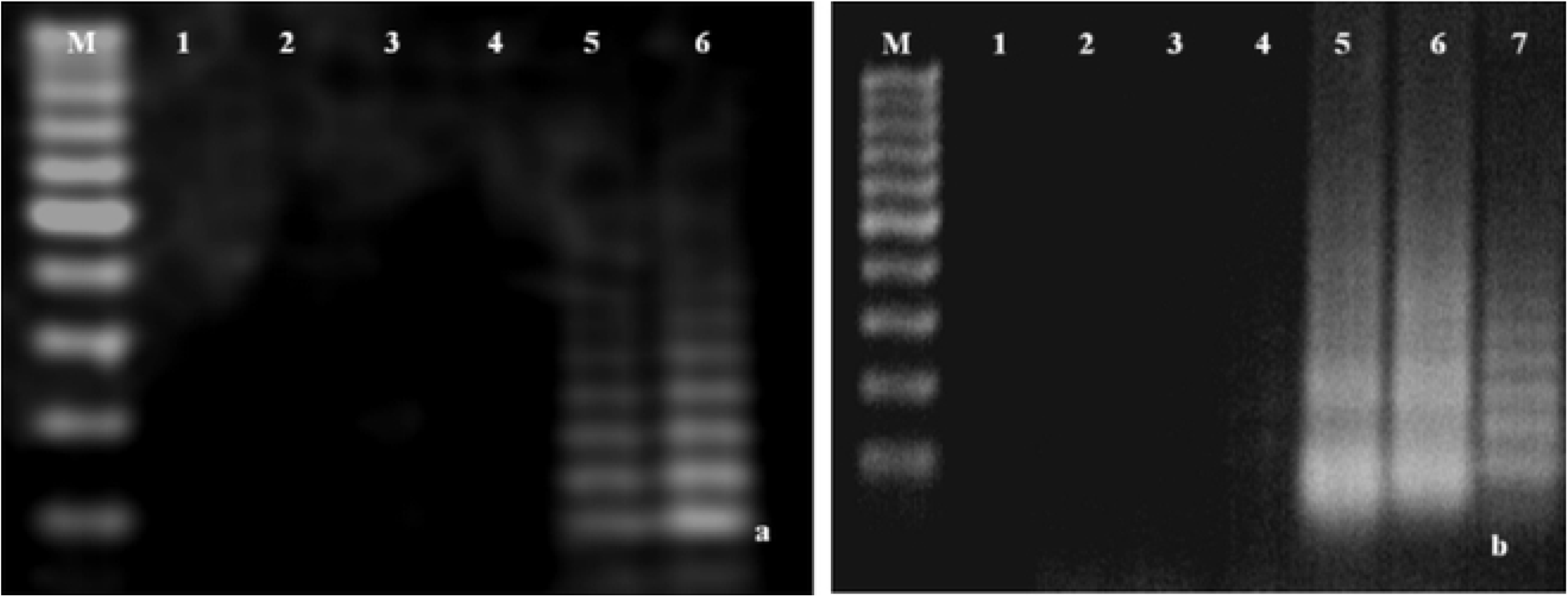
LAMP amplification of *P. angolensis* DNA (LSVM 1308) using ID1-TEF primers; a-at different temperatures; M-100bp DNA ladder, 1-60°C, 2-6l°C, 3-62°C, 4-63°C, 5-64°C, 6-65°C; b- at different incubation times; M- 100 bp ladder, 1-0 min, 2-10 min, 3-20 min, 4-30min 5-40 min, 6-50 min and 7-60 min

### 3.5 Determination of the analytical accuracy of LAMP in *P. angolensis* detection

#### 3.5.1 Assessment of the sensitivity of the ID1-ITS and IDI-TEF primer sets LAMP in the detection of *P. angolensis*

The analytical sensitivities of ID1-ITS and IDI-TEF primer sets were first compared using six serial dilutions of *P. angolensis* (LSVM 1308) gDNA including 10ng/μl, 1ng/μl, 10^−1^ ng/μl, 10^−2^ ng/μl, 10^−3^ ng/μl and 10^−4^ ng/μl). However, the IDI-TEF primer set amplified the 6^th^ DNA concentration (10^−4^ ng/μl) hence two additional dilutions (10^−5^ ng/μl and 10^−6^ ng/μl) were included to test the lowest detection limit of the IDI-TEF primer set. The detection limit of IDI-ITS primer set was 1×10^−3^ ng/μl while that of IDI-TEF primer set was 1×10^−5^ ng/μl as shown in figs 7a and b, respectively.

**Fig 7.**
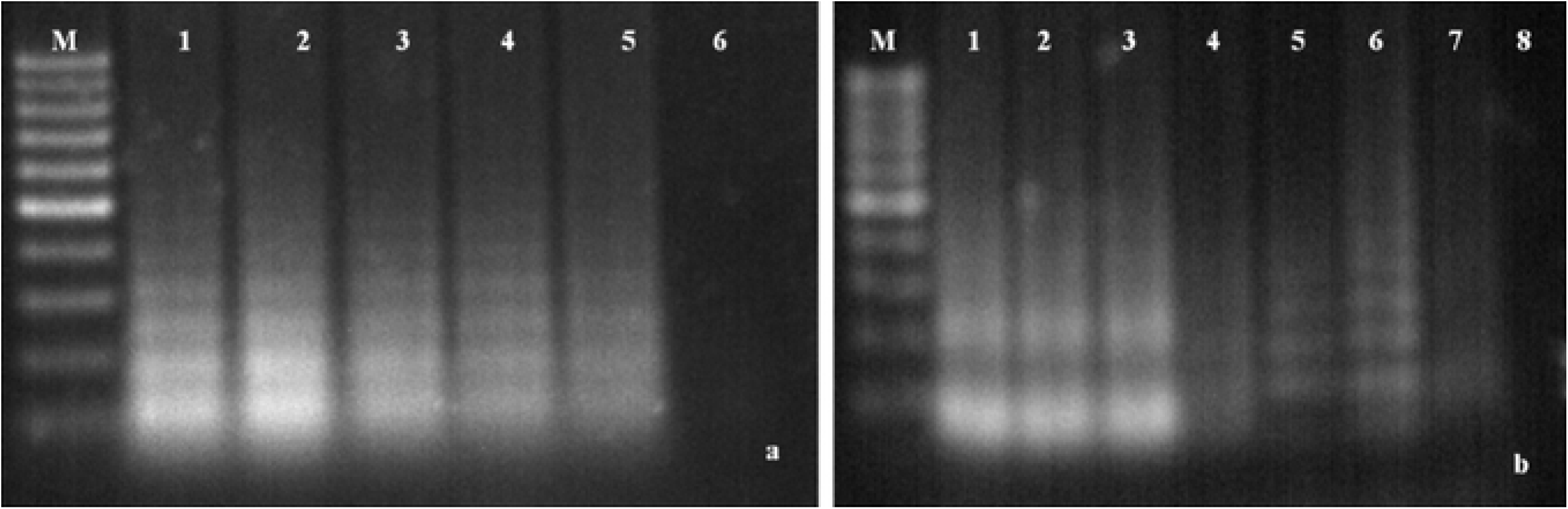
Sensitivity of ID1-ITS (a) and IDI-TEF (b) primers in *P. angolensis* detection; M-100 bp DNA ladder, 1-10ng/μl, 2-lng/μI, 3-10^−1^ ng/μl, 4-10^−2^ ng/μl, 7-10^−3^ ng/μl, 6-10^−4^ng/μl, 7-10^−5^ ng/μl and 8-10^−6^ng/μl.

#### 3.5.2 Assessment of the specificity of IDI-ITS and IDI-TEF in the detection of *P. angolensis*

Genomic DNA (1×10^−3^ng/μl) of *Pseudocercospora angolensis* and all the non-target species were detected by the ID1 primer as shown in fig 8. The IDI-ITS primer set was 100% non-specific, hence unsuitable for LAMP detection of *P. angolensis* in sweet orange. On the other hand, genomic DNA (1×10^−4^ ng/μl) of *P. angolensis* and of three of the close relatives of *P. angolensis* namely *P. musicola*, *P. griseola* and *P. fijiensis* were detected by the ID1-TEF primer set. On the other hand, DNA (1×10^−4^ ng/μl) of *P. emusae*, other citrus pathogens and *C. cinensis* were not detected by this primer set (see fig 9).

**Fig 8.**
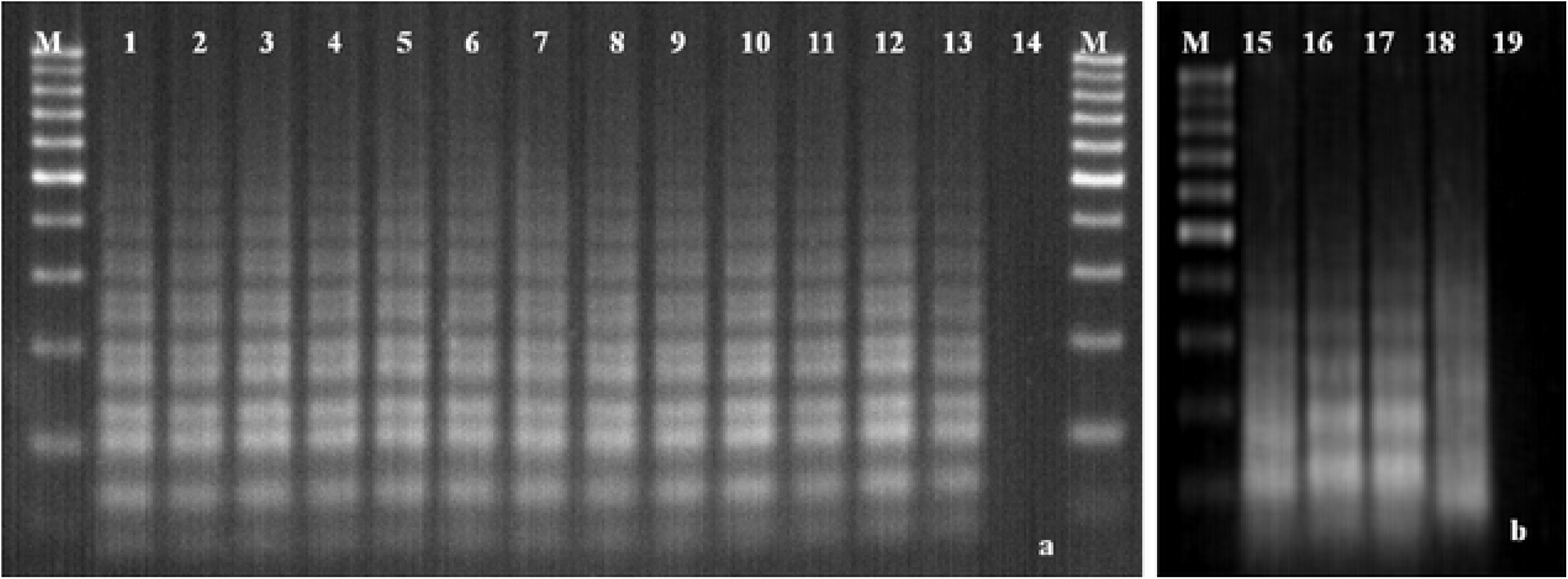
Specificity of amplification ofID1-ITS primer in *P. angolensis* detection; M-100 bp DNA ladder, 1 & 15 - Isolate of *P. angolensis*, 2- *P. musicola*, 3- *P. emusae*, 4- *P. griseo/a, 5-P fljiensis*, 6- *C. cine.sis*, 7- *C. gleosporoides*, 8- *E.fawcetii*, 9- *G, citricarpa*, 10- *A. alternata*, 11- *C. lindemuthiannum*, 12- *D. citri*, 13- *P. citrophthora*, 14&19 - Negative control (PCRgrade water),16- LSVM 09, 17- LSVM 08 and l8- MC 39.

**Fig 9.**
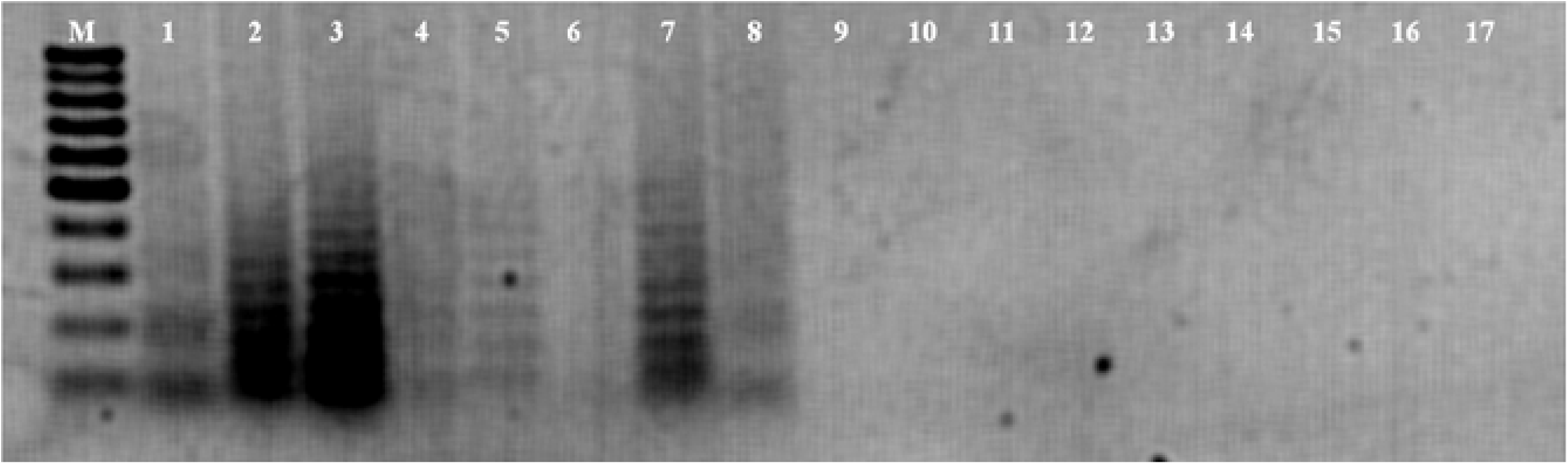
Evaluation of the specificity of the IDI-TEF primer in *P. angolensis* detection: M- 100 bp ladder, 1- Isolate of *P. angolensis*, 2- LSVM 1309, 3- LSVM 1308, 4- MC 39, 5- *P. musicola*, 6- *P. enusae*, 7- *P. griseola*, 8- *P. fijiensis*, 9- *C. cinensis*, 10- *C. gleosporoides*, 11- *E. fawcelli*, 12- *G. cilricarpa*, 13- *A. alternala*, 14- *C. lindemuthianum*, 15- *D. citri*, 16- *P. citrophthora* and 17- PCRgrade water.

Since the IDI-TEF primer was more specific than IDI-ITS primer set, the IDI-TEF primer set was chosen to assess the diagnostic accuracy of LAMP in *P. angolensis* detection.

### 3.6 Comparison of the analytical accuracy of LAMP using the TEF primers and PCR in the detection of *P. angolensis* using Cer 2 primers

Generally, the analytical specificity of both assays is sufficient for detection *P. angolensis* in sweet orange. PCR was more specific than LAMP as it detected *P. angolensis* in DNA samples known to have been extracted from *P. angolensis* which included the reference samples LSVM 1309, LSVM 1308, MC 39 and an isolate of *P. angolensis.* Other DNA from close relatives of *P. angolensis* namely *P. musicola*, P. *fijiensis*, *P. emusae* and *P. griseola* and from common citrus pathogens present in Uganda namely *C. gleosporoides*, *E. fawcetti*, *G. citricarpa*, *A. alternata*, *D. citri, P. citrophthora* and a bean pathogen known as *C. lindemuthanium* were not amplified by PCR. On the other hand, LAMP was specific to *P. angolensis* and three of its close relatives which included *P. musicola*, *P. fijiensis* and *P. griseola*. The rest of the non-target species were not detected by LAMP. The analytical sensitivity of LAMP (10^−5^ ng/μl) was much higher than that of PCR (10ng/μl).

### 3.7 Determination of the diagnostic accuracy of LAMP and PCR in the detection of *P. angolensis*

#### 3.7.1 Assessment of the diagnostic accuracy of PCR in *P. angolensis* detection

The sensitivity, specificity, positive predicive value and negative predictive value of PCR were 32.6%, 100%, 100% and 33.1% respectively. Microscopy was significantly more sensitive than PCR in *P. angolensis* detection (*X*^*2*^ = 149.26, *P*< 2.2e-16) whereas there was no difference in the specificities of microscopy and PCR in *P. angolensis* detection (*X*^*2*^ = NaN, *P* = NA).

#### 3.7.2 Assessment of the diagnostic accuracy of LAMP in *P. angolensis* detection

The sensitivity, specificity, positive predicive value and negative predictive value of LAMP were 93%, 90%, 96.5% and 81.8% respectively. Statistical analysis showed that microscopy was significantly more sensitive than LAMP in the detection of *P. angolensis* (*X*^*2*^ = 8.38, *P* = 0.0038) whereas there was no significant difference in the specificities of microscopy and LAMP in *P. angolensis* detection (*X*^*2*^ = 3.37, *P* = 0.066).

## 4. Discussion

This study aimed at developing a LAMP system for detecting *Pseudocercospora angolensis* in sweet orange.

For purposes of specific molecular diagnostics, it is important to design primers from an appropriate genetic marker(s). The region from which the primer is designed should be conserved and unique to the target species [16]. The ITS region is very useful in identifying fungi at species level hence it is the primary barcode for the fungal kingdom [22]. That explains why in this study, the ITS1, 5.8S, ITS2 and partial sequence of the large subunit ribosomal RNA in the genome of *P. angolensis* were used to design both PCR and LAMP primers. In other studies, the *Tef1-α*, ITS1 and ITS4 have been used to design PCR primers for *P. angolensis* detection [8, 9]. The *Tef1-α* region is widely used in *P. angolensis* genus phylogeny since it is highly informative at species level [23]. Hence in this study, the *Tef1-α* sequence was used to design the LAMP primers.

Loop-mediated amplification (LAMP) was fully validated in this study alongside PCR using microscopy as a gold standard. The analytical specificity of the ID1-TEF LAMP primer was not 100% specific since it could consistently detect three closely related non-target species namely *P. griseola*, *P. musicola* and *P. fijiensis* out of 12 non-target species. However, *P. griseola* is a pathogen of the common bean, a causative agent of angular leaf spot whereas *P. musicola* and *P. fijiensis* are pathogens of bananas that cause sigatoka and black leaf streak diseases respectively [24, 25]. Therefore, it is very unlikely that these pathogens infect sweet orange plants. Non-specific detection could have arisen from the fact that the *tef*-1 alfa sequence was more conserved in *P. griseola*, *P. musicola*, *P. fijienjis* and *P. angolensis* than anticipated. The 100% non-specificity of the ID1-ITS LAMP primer could also have been due to conservation of the marker sequence across both *P. angolensis* and all the non-target species. Studies done by [10, 15] have also reported non-specific detection using the ITS as a diagnostic marker. The *tef-1* alpha sequence is more suitable a diagnostic marker for *P. angolensis* than the ITS region since it is more specific. On the other hand, PCR was 100% specific in *P. angolensis* detection because the PCR primers were picked from a region more specific to *P. angolensis*. Therefore, the analytical specificity of PCR was found to be better than that of LAMP in *P*. *angolensis* detection. [26] also found PCR to be specific to detection of *Puccinia triticina*, a pathogen of wheat leaf rust. However, in their study LAMP did not detect any non-target species unlike in this study where LAMP could detect three non-target species closely related to *P. angolensis*. [7, 16] also reported LAMP to be specific to detection of *Ralstonia solanacearum* which causes wilt in over 200 plant species and Napier stunt phytoplasma, a pathogen of Napier stunt disease, respectively. The analytical specificity of LAMP could have been higher in the other studies because they used different diagnostic markers. For instance, [26] used simple repeat markers, [16] used 16S rRNA gene whereas [7] used *fliC* sequence of *R. solanacearum*. Both LAMP and PCR were incapable of detecting sweet orange DNA, which makes them suitable for detecting *P. angolensis* in sweet orange leaf and fruit samples directly without having to first culture the pathogen. The analytical sensitivity of LAMP was higher with the ID1-TEF primer set (10^−5^ ng/μl) than with the ID1-ITS primer set (10^−3^ ng/μl) probably because the ID1-TEF primer set was more efficient in binding to the template than the ID1-ITS primer set. It could also be that the *tef* 1 alpha gene copy number per cell was higher than that of the ITS. The analytical sensitivity of LAMP (10^−5^ ng/μl) was much higher than that of PCR (10ng/μl) in *P. angolensis* detection. Previous studies conducted by [27, 26] also found LAMP to be more sensitive than PCR in the detection of bacterial meningitis and fungal wheat leaf rust, respectively. This translates to LAMP being more suitable in detecting *P. angolensis* during early pathogenesis than PCR, hence enhancing early disease control. The analytical sensitivity of LAMP was higher than that of PCR probably because the sequence that the LAMP primers were targeting (*tef-1* alpha gene) had more copies in the *P. angolensis* genome the sequence targeted by the PCR primers.

In this study, LAMP could detect *P. angolensis* in 40 minutes unlike PCR which took 1h 38min. [28] also reported LAMP to have detected *Trypanasoma cruzi* in 40 minutes. Loop mediated isothermal amplification (LAMP) produces faster results than PCR because in LAMP reactions, three primers pairs were used to generate amplicons whereas in PCR, only one primer pair was used. Reactions for LAMP use more primer pairs and it is auto cycled whereas PCR reaction cycles (denaturation, annealing and extension) are controlled at different temperatures and durations [18]. Hence LAMP has a lower turnaround time than PCR in *P. angolensis* detection, which makes LAMP more suitable for screening large numbers of sweet orange samples for *P. angolensis* than PCR.

Evaluation of the diagnostic accuracy of PCR revealed that the diagnostic sensitivity of microscopy (100%) was significantly higher than that of PCR (32.6%) (*X*^*2*^ = 149.26, *P* < 2.2e-16). This could have been because PCR detection of *P. angolensis* in the samples depended on the DNA concentration whereas microscopy depended on the presence or absence of *P. angolensis* conidia in cultures isolated from the leaf samples and not concentration thereof. Given that the DNA was obtained from lesions caused by *P. angolensis* on the sweet orange leaf samples, the amount of *P. angolensis* or spores thereof could have been too little to achieve the detection limit of PCR during DNA extraction. On the other hand, there was no difference in the specificities of PCR and microscopy since they both did not detect *P. angolensis* in the sweet orange leaf samples asymptomatic for *Pseudocercospora* leaf and fruit spot disease. The diagnostic specificity was excellent for PCR because the primers were not complementary to citrus DNA which was present in the samples asymptomatic for *Pseudocercospora* leaf and fruit spot disease. The positive predictive value of PCR was excellent (100%) whereas the negative predictive value was low (33.1%). Therefore, PCR has 100% chance of correctly identifying truly positive samples and 33.1% of identifying negative samples in the diagnosis of *P. angolensis* [29]. The high positive predictive value was due zero (0) false positives which could have been because of the perfect analytical specificity of PCR in detection of *P. angolensis* whereas the low negative predictive value was because PCR had very many false negatives (101/150). The high false negative rate could have been because PCR reactions can easily be inhibited by biological substances in the sample [30]. It could also be that the DNA concentration of *P. angolensis* extracted from the leaf lesions was too low to be detected by PCR. The low negative predictive value makes PCR unattractive for screening sweet orange leaf/fruit samples for *Pseudocercospora* leaf and fruit spot disease for disease control since it would falsely classify most of the diseased citrus plants as disease-free.

There was a significant difference between the diagnostic sensitivities of LAMP (93%) and microscopy (100%) (*X*^*2*^ = 8.38, *P* = 0.0038). However, the diagnostic specificities of LAMP (90%) and microscopy (100%) were not significantly different (*X*^*2*^ = 3.37, *P* = 0.066). The significant difference in the sensitivities of LAMP and microscopy is not surprising since their principle of detection differs whereby LAMP uses *P. angolensis* DNA whereas microscopy uses *P. angolensis* conidia. The diagnostic sensitivity of LAMP was as high as 93% because LAMP was able to detect very little concentrations of DNA extracted from small quantities of *P. angolensis* and spores thereof on leaf lesions. On the other hand, the diagnostic specificity of LAMP was as high as 90% because six primers were used to amplify the target sequence in the genome of *P. angolensis* DNA. The high positive predictive value (96.5%) and negative predictive value (81.8%) of LAMP could have been because LAMP reactions are tolerant to sample matrix inhibitors^15^ which is likely to contribute to the low false negative rate and false positive rates, respectively. Our results were similar to those reported by [31] who reported high sensitivity, specificity, PPV and NPV of 98.3%, 100%, 100% and 98%, respectively in LAMP detection of malaria, using microscopy as the gold standard. The negative predictive value of LAMP makes it an attractive tool for disease management because only sweet orange seedlings/fruits free of *P. angolensis* should be certified for sale or planting. Since the positive predictive value of LAMP is very high, it also enhances identification of diseased plants for disease control through application of fungicides and removal of diseased plant parts.

## 5. Conclusion

The LAMP assay developed in this study can be applied in faster and early diagnosis of *Pseudocercospora angolensis* in sweet orange plants. Early diagnosis will enhance timely management of *Pseudocercospora* leaf and fruit spot disease.

## Acknowledgements

This study was supported with funding from the Korea Program on International Agriculture (KOPIA) of the Rural Development Administration (RDA), Republic of South Korea. We acknowledge Allan Male and Catherine Acam of CIAT-Uganda for their guidance during laboratory analyses, and Jaqueline Hubert ANSES- France for availing the reference DNA samples and microorganisms.

